# Coronaviruses use ACE2 monomers as entry receptors

**DOI:** 10.1101/2023.01.25.525479

**Authors:** Patrick Eiring, Teresa Klein, Simone Backes, Marcel Streit, Sören Doose, Gerti Beliu, Markus Sauer

## Abstract

The angiotensin-converting enzyme 2 (ACE2) has been identified as entry receptor on cells enabling binding and infection with the severe acute respiratory syndrome coronavirus 2 (SARS-CoV-2) via trimeric spike (S) proteins protruding from the viral surface^1,2^. It has been suggested that trimeric S proteins preferably bind to plasma membrane areas with high concentrations of preferably multimeric ACE2 receptors to achieve a higher binding and infection efficiency^1,3^. However, our current knowledge about the influence of ACE2 expression and organization in the plasma membrane on SARS-CoV-2 infection efficiency remains elusive. Here we used *direct* stochastic optical reconstruction microscopy (*d*STORM) in combination with different labeling approaches to visualize the distribution and quantify the expression of ACE2 on different cells. Our results reveal that endogenous ACE2 receptors are present as monomers in the plasma membrane with densities of only 1-2 receptors μm^-2^. In addition, binding of trimeric S proteins does not induce clustering of ACE2 receptors in the plasma membrane. Supported by infection studies using vesicular stomatitis virus (VSV) particles bearing S proteins our data demonstrate that a single S protein interaction per virus particle with a monomeric ACE2 receptor is sufficient for infection which attests SARS-CoV-2 a high infectivity.

## Introduction

Severe acute respiratory syndrome coronavirus 2 (SARS-CoV-2) is the third human coronavirus that recently emerged. It is spreading rapidly in humans, causing COVID-19, respiratory syndromes with severe and often fatal progression^4–7^. The SARS-CoV-2 genome shares ~80% identity with that of SARS-CoV responsible for the SARS pandemic in 2002^8^. All human coronaviruses, including SARS-CoV, MERS-CoV and, SARS-CoV-2 are enveloped, single-stranded, positive-sense RNA viruses. Each SARS-CoV-2 virus particle (virion) is equipped with 24-40 homotrimeric transmembrane spike (S) glycoproteins that are key to binding and fusing with human cells^9^. After more than two years of investigations, it is undisputed that SARS-CoV-2 invades human cells by binding with the S1 subunit via the receptor-binding domain (RBD) to the peptidase domain of angiotensin-converting enzyme 2 (ACE2) as entry receptor^1^. Through binding to ACE2, another cleavage site on subunit S2 is exposed, followed by spike priming and fusion using the cellular serine protease TMPRSS2^2,10^. At low protease levels, entry via endosomal uptake can also occur^11^. The fact that antiserum raised against human ACE2 completely blocks infection with SARS-CoV-2 demonstrates that binding of the virus to the receptor is the decisive first step in infection^2^.

Since the S protein undergoes large conformational changes during the fusion process, it must be highly flexible and dynamic. Especially the RBD in the S1 subunit was revealed to be considerably flexible, exhibiting two conformational states, an inactive *down* state and an active *up* state, whereby only the *up* state can bind to ACE2. Recently, it has been shown that the trimeric structure of the S protein of SARS-CoV-2 exists with one RBD in an *up* and two in *down* conformations^12^ suggesting stoichiometric binding of one S protein per ACE2 receptor. Furthermore, high flexibility of the spikes themselves allows them to sway and rotate, thus possibly enabling multiple spikes per virus to bind to a human cell^3^. This, however, would require a high expression level of ACE2 or the occurrence of ACE2 oligomers or preformed clusters in the plasma membrane to increase the chance for neighboring spikes to find binding sites. Indeed, high-resolution cryo-electron microscopy structures obtained on the recombinant full-length human ACE2 in contact with the RBD suggest that ACE2 forms homodimers whereby each monomer can bind a S-glycoprotein trimer^1^.

These findings are of utmost importance for the structure-based rational design of binders with enhanced affinities to either ACE2 or the S protein of coronaviruses. However, structural investigations of SARS-CoV-2 – ACE2 interactions have been performed with purified proteins and thus do not reflect the native environment, i.e., the plasma membrane of cells, with its particular organization principles. For example, it has been shown that viruses preferentially bind to detergent-resistant ordered plasma membrane domains, i.e., glycolipid nano- and microdomains, to penetrate the cell, e.g., because of the enrichment of their receptors in these domains^13,14^. Hence, it has to be investigated if ACE2 exists in preformed dimers or oligomers in the plasma membrane and, whether these clustering sites are enriched in glycolipid domains that are important for invagination of the membrane and endocytosis of the viral particle^1,11,15^. In addition, ACE2 expression has been investigated only indirectly by quantifying RNA expression in nasopharyngeal and bronchial samples or using immunoblotting with polyclonal anti-ACE2 antibodies^16^. For example, in one of the investigations, it has been found that decreased expression of ACE2 is associated with cardiovascular diseases^17^. On the other hand, patients with COVID-19 displayed an average three-fold increase in ACE2 expression that may promote multi-organ failure^18,19^. In addition, a lower expression of alveolar ACE2 in young children compared to adults has been associated with a lower prevalence of COVID-19 in children^20^. However, protein expression levels in cells are often poorly predicted by mRNA-transcript levels^21^.

Despite the outstanding importance for SARS-CoV-2 drug design, knowledge about the expression level of ACE2 receptors in the plasma membrane of target cells and their interactions with trimeric spike proteins remains elusive. Here, we have set out to visualize the distribution, investigate their oligomeric state and quantify the number of endogenous ACE2 receptors in the plasma membrane of Vero, Vero E6, U2-OS, COS-7, HEK293T, and ACE2 overexpressing HEK293T cells by single-molecule sensitive super-resolution fluorescence imaging using fluorescently labeled primary antibodies. The influence of ACE2 expression level on infectivity was verified using replication-defective vesicular stomatitis virus (VSV) particles bearing coronavirus S proteins^22^. Our results demonstrate that endogenous ACE2 is present as a monomer in the plasma membrane in the absence and presence of S protein and does not form dimers or higher aggregates. Furthermore, our data imply that the infection efficiency is also determined by other factors, including lipid composition of the plasma membrane and possibly other molecules among the 332 interaction partners so far identified^23^.

## Results

### Imaging and quantification of plasma membrane ACE2 receptors by *d*STORM

It is common ground that binding of the S protein to the ACE2 receptor is a critical step for SARS-CoV-2 to infiltrate target cells. Infection studies using replication-defective vesicular stomatitis virus (VSV) particles bearing coronavirus S proteins^22^ revealed that most human cell lines are susceptible to infection^2^. Therefore, we selected Vero (African green monkey kidney cells), Vero E6, and HEK293T cells (a transfectable derivative of human embryonic kidney 293 cells) as accepted reference systems for SARS-CoV research in cell-culture-based infection models^2,24,25^. In addition, we overexpressed ACE2 in HEK293T cells to investigate the influence of higher ACE2 expression levels on infection efficiency. Furthermore, we used COS-7 (a derivative of the African green monkey kidney fibroblast cell line CV-1) and U2-OS as negative controls since fibroblast cell lines have shown negligible infection efficiency^2,24^, which has been attributed to the low ACE2 expression level^26^.

For fluorescence imaging and quantification of endogenous plasma membrane ACE2, we used single-molecule sensitive super-resolution imaging by *d*STORM^27^. To extract quantitative information from single-molecule localization data, the average number of blinking events measured for a single fluorescently labeled probe (e.g. antibody) can be used to determine the number of bound probes, which corresponds to the expression level of accessible endogenous plasma membrane proteins^28,29^. We tested different monoclonal antibodies with respect to binding specificity and selected the monoclonal anti-ACE2 antibody from Biolegend (clone: A20069I), which showed the lowest non-specific binding tendency. To facilitate quantification, we labeled the primary antibody with Alexa Fluor 647 (AF647) at a degree of labeling (DOL) of 2.5. Cells were fixed with 4% methanol-free formaldehyde and stained for ACE2 receptors with primary antibody for 1 h in PBS. After post-fixation with 4% formaldehyde / 0.2% glutaraldehyde, *d*STORM imaging of cells was performed by total internal reflection fluorescence (TIRF) microscopy to selectively image the basal plasma membrane at a high signal-to-noise ratio (Fig. 1a). To ensure saturation of accessible antigen epitopes on the plasma membrane we titrated the antibody concentration in separate experiments and used antibody concentrations of 4 μg ml^-1^ in all *d*STORM quantification experiments (Supplementary Fig. 1).

**Fig. 1.**
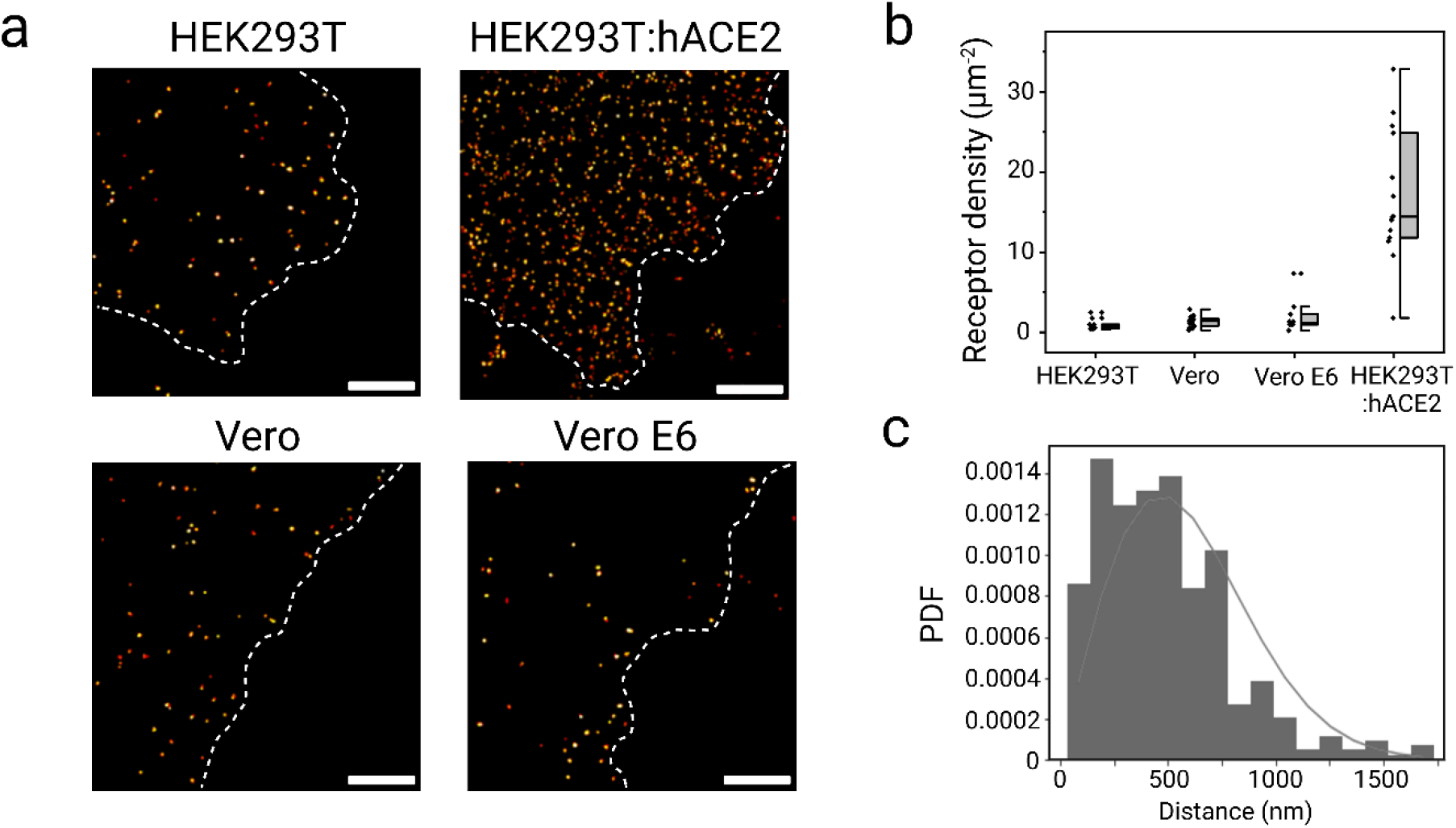
*d*STORM imaging and quantification of ACE2 on different cell lines. **a,** *d*STORM example images of HEK293T, ACE2-overexpressing HEK293T (HEK293T:hACE2), Vero, and Vero E6 cells indicating a homogeneous distribution of ACE2 receptors in the basal plasma membrane. **b**, Boxplots show localization cluster densities translated into ACE2 receptors μm^-2^ of 1.0 ± 0.2 (HEK293T), 17.0 ± 2.2 (HEK293:hACE2), 1.4 ± 0.1 (Vero) and 2.0 ± 0.7 (Vero E6) (n=10-30 cells). **C**, Probability density function (PDF) of ACE2 distances in the plasma membrane of Vero E6 cells showing that the average distance between ACE2 receptors is ~500 nm. The gray line represents the theoretical PDF of receptors homogeneously distributed in 2D. Scale bar, 3 μm (a).

*d*STORM images revealed a homogeneous distribution of ACE2 in the plasma membrane of Vero, Vero E6, HEK293T, and ACE2 overexpressing HEK293T cells (Fig. 1a and Supplementary Fig. 2). To study the distribution of plasma membrane ACE2 receptors, we used a density-based spatial clustering of applications with noise (DBSCAN) algorithm with customized localization analysis (LOCAN)^30^. After selecting basal membrane regions that do not show folded membrane areas, repeated localizations were grouped using the DBSCAN algorithm with appropriate parameters to ensure each cluster represents an isolated receptor^31^. The Ripley-h function confirms that localization clusters only occur on the length scale of the localization precision (Extended Data Fig. 1a). Under the applied experimental and dilute labeling conditions, we localized each AF647 antibody on average 9.2 ± 0.2 times. Hence, each localization cluster with on average ~9 localizations corresponds to a labeled ACE2 receptor. The overall density of ACE2 in the plasma membrane was low enough to yield well-separated nearest neighbors for all plasma membranes investigated (Fig. 1a). In each experiment, 10-20 cells were analyzed to obtain ACE2 density distributions. For both Vero cell lines similar amounts of ACE2 receptors could be identified with 1.4 ± 0.1 (S.E.) and 2.0 ± 0.7 (S.E.) receptors μm^-2^ in Vero and Vero E6, respectively (Fig. 1b). ACE2 levels were slightly lower for HEK293T cells with 1.0 ± 0.2 (S.E.) receptors μm^-2^. Transfection of HEK293T cells led to overexpression of ACE2 receptors with densities of 17.0 ± 2.2 (S.E.) ACE2 receptors μm^-2^. Both COS-7 and U2-OS control cell lines showed negligible ACE2 densities well below 0.5 receptors μm^-2^, most probably due to the non-specific binding of antibodies on the membrane (Supplementary Fig. 3). These results show that endogenous ACE2 receptors are expressed homogenously but at low densities of ≤ 2 μm^-2^ in the plasma membrane of Vero cells (Fig. **1b**), which translates into average distances of ACE2 receptors of ~500 nm (Fig. 1c).

### ACE2 expression levels correlate with VSV infection efficiency

To investigate if the ACE2 expression directly determines the infection efficiency, the different cells lines were infected with VSV particles bearing coronavirus S proteins (VSV-S^GFP^)^2^. In the genome of VSV-S^GFP^ particles, the coding sequence of the endogenous surface protein G is deleted and replaced with the GFP open reading frame. By infecting S protein expressing cells, newly budding particles contain the S protein as the only surface protein. Upon infection with VSV-S^GFP^, cells express green fluorescent protein (GFP). To measure the percentage of infected cells, nuclei were stained with DAPI (Fig. 2a). Albeit ACE2 expression levels are similar on both Vero cell lines, infection with VSVs was only possible for Vero E6 but not for Vero cells, i.e., 6.1% ± 4.1% (S.D.) of Vero E6 cells showed GFP expression whereas Vero as well as COS-7 and U2-OS did not show GFP signal after 24 h of infection (Fig. 2b and Supplementary Fig. 4). Interestingly, the infection efficiency of Vero E6 cells was substantially (~5-fold) higher than observed for HEK293T cells (Fig. 2b), although the ACE2 expression levels differ only by a factor of 2 (Fig. 1b). Overexpressing ACE2 HEK293T cells showed the highest infection efficiency of 28.4% ± 12.6% (S.D.) (Fig. 2b) in accordance with the higher ACE2 expression level (Fig. 1b).

**Fig. 2.**
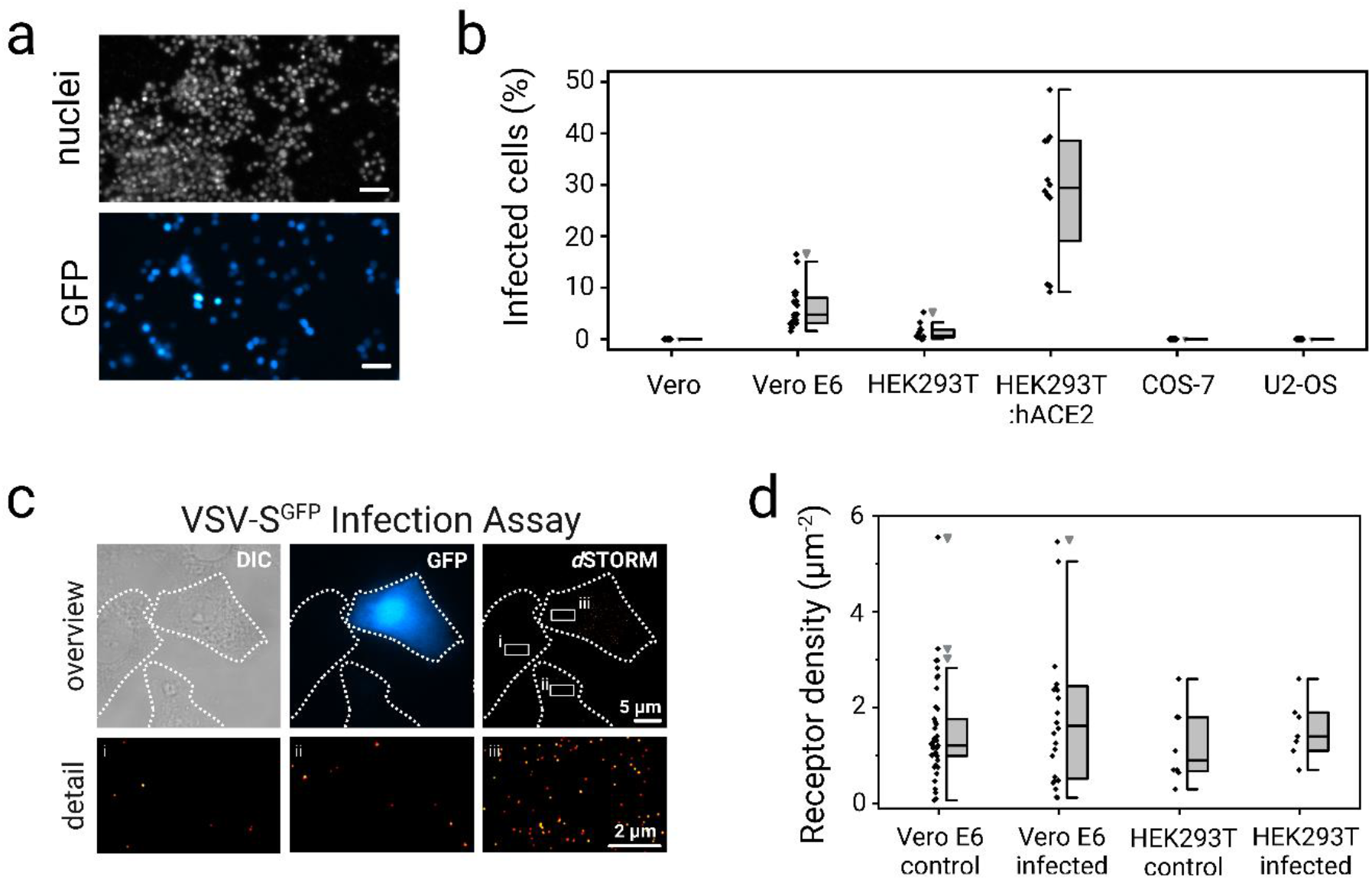
Infectivity of cells does not solely depend on ACE2 expression. **a**, Widefield fluorescence images of GFP and DAPI of ACE2 overexpressing HEK293T cells after 24 h infection with VSV-S^GFP^. **b**, Percentage of cells infected (showing GFP signal) after 24 h infection with VSV-S^GFP^. **c**, VSV-S^GFP^ infection assay comparing infection (widefield GFP signal) and ACE2 expression (*d*STORM) in Vero E6 cells. Small panels (i-iii) display magnifications of boxed regions in the *d*STORM image. **d**, ACE2 receptor density (μm^-2^) determined by *d*STORM in infected and uninfected Vero E6 and HEK293T cells. Scale bar, 50 μm (a), 5 μm (c), and 2 μm (magnified views (c)).

To examine if predominantly those cells were infected that express higher ACE2 levels, we quantified and compared ACE2 expression levels on infected and non-infected cells in the same experiment. Therefore, cells were fixed after 24 h incubation with VSV-S^GFP^ particles and immunostained for ACE2. *d*STORM was performed on cells showing GFP signal (infected) and cells without GFP signal (uninfected) within the same well (Fig. 2c). Here, infected Vero E6 and HEK293T cells showed only slightly higher ACE2 expression levels (Fig. 2d). These results demonstrate that ACE2 receptors are necessary for SARS-CoV-2 infection and infection efficiency correlates with ACE2 expression for the same cell line. However, the infection efficiency can differ strongly for different cell lines indicating that other factors play at least a supporting role in infection.

### Oligomeric state of ACE2 in the plasma membrane

The stoichiometry of plasma membrane proteins is an important determinant of function and interactions between individual proteins. Previous reports suggested the binding of multiple spikes per virus to ACE2 homodimers or oligomers in the plasma membrane via stoichiometric binding of one S protein per ACE2 receptor^1,3,12^. Therefore, it is easy to believe that ACE2 receptors are prearranged as multimers in the plasma membrane, or, alternatively, form multimers upon binding of S-glycoprotein trimers to increase the binding strength and infection efficiency. To verify the existence of receptor multimers in the plasma membrane of cells, photoactivated localization microscopy (PALM) in combination with photoactivatable fluorescent proteins such as mEos fused to the protein of interest has been the method of choice^32,33^. However, visualization of endogenous proteins requires immunolabeling and thus remains more challenging.

To investigate the capability of *d*STORM to distinguish endogenous monomeric from dimeric plasma membrane proteins, we imaged monomeric CD18^34^, homodimeric CD69^35^ as well as heterodimeric CD11a/CD18^34^ by *d*STORM on Jurkat cells using primary AF647 labeled antibodies for immunostaining (Figs. 3a-c). Analysis of *d*STORM data revealed mean localization numbers per localization cluster of 7.3 ± 0.2 (S.E.) for monomeric CD18, 14.2 ± 0.3 (S.E.) for dimeric CD69 and 16.3 ± 0.4 (S.E.) for heterodimeric CD11a/CD18. The dimeric receptor population is indicated by the shift towards higher localization numbers per cluster in the probability density function (PDF) of localizations detected per spatially separated fluorescent signal (localization cluster) of CD69 and CD11a/CD18 (Fig. 3d).

**Fig. 3.**
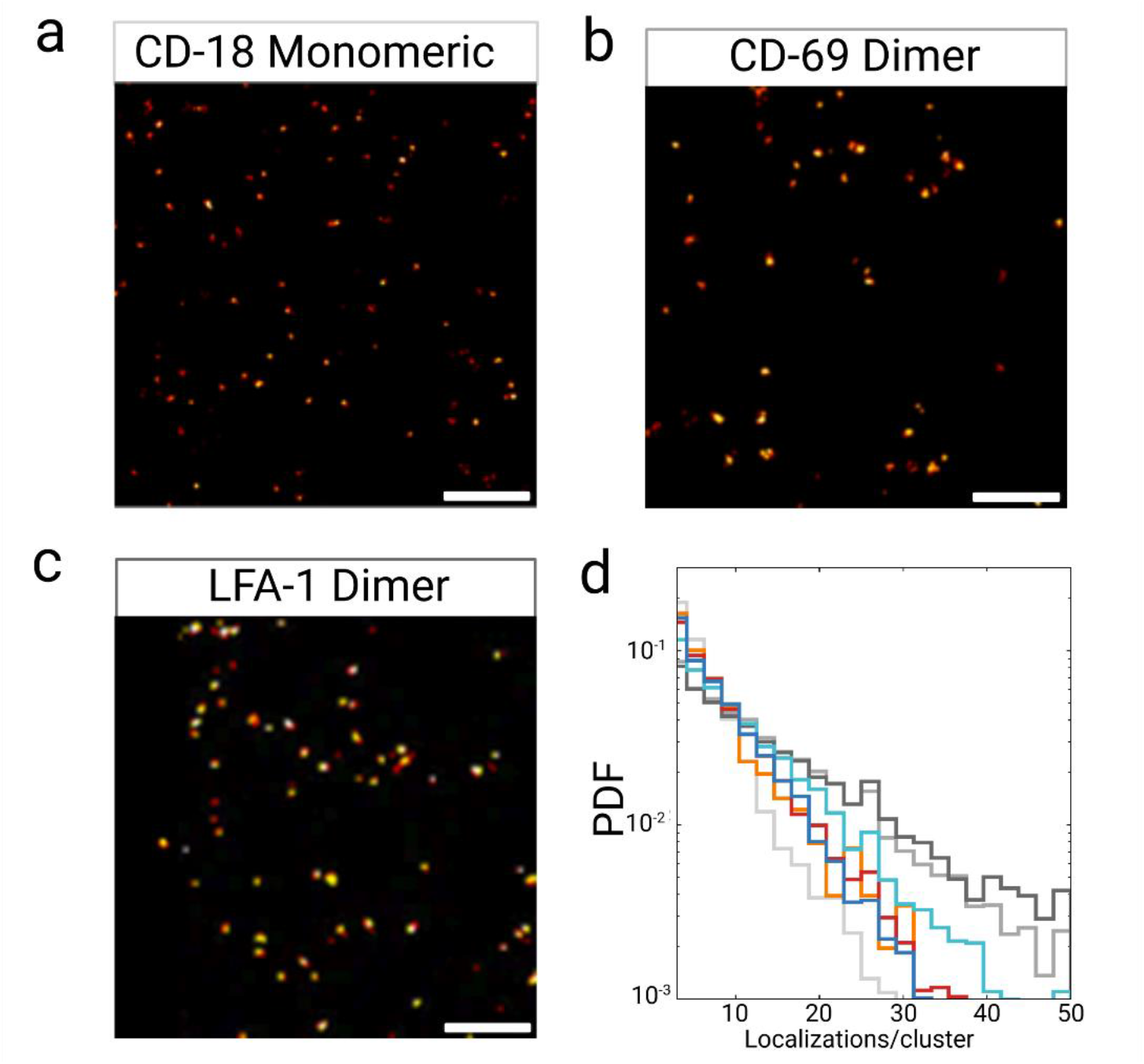
*d*STORM with antibodies reveals the oligomeric state of receptors. *d*STORM images of monomeric CD18^33^ **(a)**, homodimeric CD69^34^ **(b)** as well as heterodimeric LFA-1^33^ **(c)** on Jurkat cells using primary labeled antibodies for immunostaining. **d**, PDFs of the localization numbers detected for the three receptors CD18 (light grey), CD69 (grey), and LFA-1 (dark grey) showing a broader distribution towards higher localization numbers per spatially isolated localization cluster for dimeric receptors. PDFs of the localization numbers detected for ACE2 on HEK293T (orange), on HEK293T:ACE2 (red), Vero E6 (cyan) or Vero (blue) demonstrate the organization of ACE2 as monomers in the plasma membrane. Scale bar, 3 μm.

*d*STORM imaging performed with primary AF647 labeled anti-ACE2 antibodies showed similar results as detected for CD18. The PDF of localization numbers showed no indication of dimer or higher aggregate formation, demonstrating the presence of monomeric ACE2 receptors in the plasma membrane of Vero, Vero E6, and HEK293T cells (Fig. 3d). Even ACE2 overexpressing HEK293T (HEK293:hACE2) cells demonstrate the presence of ACE2 as a monomer in the plasma membrane (Fig. 3d). To confirm that endogenous dimeric receptors can indeed be identified by *d*STORM of adherent HEK293T cells we performed immunostaining of Neuropilin-1, a dimeric receptor important for SARS-CoV-2 entry and infectivity^36^. *d*STORM imaging and analysis using anti-NRP-1 antibodies confirmed the existence of dimeric NRP-1 receptors on HEK293T cells (Extended Data Fig. 2a). The PDF of localization numbers for NRP-1 localization clusters (Extended Data Fig. 2b) is similar to PDFs recorded for dimeric CD69 and heterodimeric CD11a/CD18 (Fig. 3d). Interestingly, HEK293T cells express ~3 NRP-1 μm^-2^, but Vero as well as Vero E6 cells do express NRP-1 only at negligible levels in the plasma membrane (Extended Data Fig. 2c) indicating a less important role of NRP-1 for SARS-CoV-2 infections in Vero cells.

Finally, we used click labeling of genetically modified ACE2 receptors with tetrazine-dyes to exclude that the detection of monomeric ACE2 is caused by the size of IgG antibodies and related steric hindrance preventing labeling of dimeric receptors with two antibodies. For site-specific incorporation of the non-canonical amino acid (ncAA) TCO*-L-lysine into human ACE2 (pCG1-hACE2)^37,38^ in COS-7 cells, we used genetic code expansion (GCE)^39,40^. The method enables site-specific efficient labeling of intra- and extracellular proteins with minimal linkage error with organic tetrazine-dyes via bioorthogonal click labeling^41^ while preserving the functionality of proteins^42,43^. The generated click mutant (pCG1-ACE2^N137TAG^) was tested for ncAA incorporation and click labeling. Here, the mutant N137TAG showed efficient labeling demonstrated by a continuous fluorescence signal visible along the cell membrane (Supplementary Fig. 5). After click labeling of ACE2 in living COS-7 cells with tetrazine-AF647 (Tet-AF647), they were either directly fixed or first treated with recombinant trimeric spike protein before fixation, followed by *d*STORM imaging (Extended ata Fig. 1c). We computed nearly identical Ripley-h functions from all recorded localizations with and without spike addition. Comparison with synthetic data that resembles *d*STORM blinking conditions and that clusters homogenously distributed on identical region of interest suggests no further clustering in the experimental data. Also, DBSCAN cluster analysis revealed monomeric ACE2 distributions and no alteration upon spike addition (Extended Data Figs. 1a,b). Localization data analysis thus demonstrated that the treatment with trimeric spike protein does not induce the formation of ACE2 multimers corroborating that ACE2 is expressed as a monomer in the plasma membrane that do not multimerize upon binding of trimeric spike proteins^1^.

## Discussion

The ACE2 receptor is of fundamental importance for infections with SARS-CoV-2, but knowledge about its expression level, distribution, and oligomeric state in the plasma membrane of target cells has remained elusive. Here we used immunolabeling with primary monoclonal antibodies and super-resolution fluorescence imaging by *d*STORM to visualize the distribution and quantify the expression of endogenous ACE2 receptors in the plasma membrane of various cell lines. Our results show that endogenous ACE2 is homogenously expressed in the plasma membrane of HEK293T, Vero, and Vero E6 cells that are accepted cell-culture-based infection models for SARS-CoV-2 infection^2,25,26^ with expression levels of 1-2 ACE2 molecules μm^-2^ (Fig. 1a,b and Supplementary Fig. 2). As expected, on the plasma membrane of cells like COS-7 and U2-OS that are not subjected to SARS-CoV-2 infections, ACE2 expression is negligible (Supplementary Fig. 3).

Furthermore, infection studies with VSV-S^GFP^ particles in combination with *d*STORM imaging corroborated the finding that ACE2 represents the key receptor for SARS-CoV-2 entry, i.e. infected cells generally expressed slightly more ACE2 molecules on their surface (Fig. 2). However, our data show that the infection efficiency differs strongly between different cell lines. A striking example represents the different infection efficiency of Vero and Vero E6 cells (Fig. 2b). While they exhibit similar ACE2 expression levels (Fig. 1a,b), Vero cells are not infected by VSV-S^GFP^ particles. Therefore, we hypothesized that other co-receptors of SARS-CoV-2 entry, such as NRP-1^36^, are expressed at substantially higher concentrations on Vero E6. However, our data demonstrate that NRP-1 does not play a role in the infectivity of Vero cells (Extended Data Fig. 2).

Since glycosphingolipids are important host cell targets for different pathogens and are supposed to represent endocytotic entry sites^14^, we investigated the impact of sphingolipids on infectivity. In this context, glycosphingolipids such as monosialotetrahexosylganglioside GM1, a prototype ganglioside, which interacts with protein receptors within lipid rafts to generate signaling platforms are important^44,45^. Therefore, we visualized the lipid raft marker GM1 by staining with AF647 labeled cholera toxin subunit B. Fluorescence imaging revealed a high abundance of GM1 in the plasma membrane of Vero E6 and a lower concentration in Vero cells (Extended Data Fig. 3), indicating that glycosphingolipids support viral entry via ACE2 receptors^46^. COS-7 cells also exhibit a high GM1 concentration in their plasma membranes (Extended Data Fig. 3) but cannot be infected by SARS-CoV-2^2,25^, which is due to the negligible small expression of ACE2 in the plasma membrane (Supplementary Fig. 3). Overall, these data demonstrate that ACE2 receptors are necessary but not sufficient for SARS-CoV-2 infection.

The so far unanswered question of SARS-CoV-2 infection that still remained is if pre-clustering or binding-induced clustering of ACE2 receptors in dimers, trimers, or higher aggregates is crucial for viral entry. Intuitively one expects that a trimeric S protein can bind to several ACE2 receptors in the plasma membrane of target cells to improve binding strength and thus the infection probability^1,12,13^. These considerations are in line with the finding that higher ACE2 expression levels promote infection with SARS-CoV-2 (Fig. 1b and 2d)^18–20^. To address this question, we performed *d*STORM experiments with reference receptors that form monomers, dimers, and heterodimers in the plasma membrane using AF647 labeled primary antibodies. The refined analysis of localization numbers per localization cluster detected revealed that *d*STORM with primary antibodies can distinguish between endogenous monomeric and dimeric receptors and showed that ACE2 is present as a monomer in the plasma membrane (Fig. 3). Binding-induced clustering was investigated by introducing an unnatural amino acid into ACE2 at a site where it does not interfere with RBD binding followed by click labeling with Tet-AF647 and *d*STORM imaging before and after addition of trimeric S proteins. The obtained data clearly revealed that binding of one S protein subunit to one ACE2 receptor does not induce clustering to improve the binding strength (Extended Data Fig. 1). Considering the average distance between neighboring ACE2 receptors of ~500 nm (Fig. 1c) supports this finding because it appears unlikely that binding of a trimeric S protein to a monomeric ACE2 receptor can unite other ACE2 receptors in reasonable time that are separated at such large distances. Furthermore, such low ACE2 densities make it unlikely that more than one S protein per virus particle can interact with a cell^3^. Thus, it appears that electron microscopy structures obtained from interacting recombinant expressed proteins do not necessarily reflect interactions of plasma membrane receptors embedded in their native environment with viral proteins. To conclude, our data unequivocally demonstrate that the molecular mechanisms of SARS-CoV-2 infection are not understood in detail. Yet, we hope that the presented quantitative molecular understanding of the ACE2 – S protein interaction on the cell membrane offers perspectives for the development of improved drugs for the treatment of SARS-CoV-2 infections.

## Acknowledgments

The authors thank E. Maier for cell culture support. The expression plasmid of hACE (pCG1-hACE2) was a gift to S.B. from M. Hoffmann (Göttingen, Germany). The plasmid for the expression of the tRNA/aminoacyl transferase pair (pNEU-hMbPylRS-4xU6M15, herein termed (PylRS/tRNA^Pyl^) was a gift from I. Coin (Addgene, no. 105830). P.E., T.K. and M.S. received funding from the European Research Council (ERC) under the European Union’s Horizon 2020 research and innovation programme (grant agreement No 835102), the Deutsche Forschungsgemeinschaft (DFG SA829/19-1) and the Bundesministerium für Bildung und Forschung (BMBF, grant #13N15986).

## Author contributions

G.B. and M.S. conceived, designed and supervised the project. P.E. and T.K. performed all *d*STORM, widefield and quantification experiments. S.B. produced VSV particles. S.B. and T.K. performed infection experiments. G.B. and M.St. performed ncAAs incorporation and click-labeling. S.D. performed data analysis. G.B. and M.S. wrote the manuscript. All authors revised the final manuscript.

## Competing interest

The authors declare no competing interests.

## Data availability

The data that support the findings of this study will be provided by the corresponding author upon reasonable request.

## Code availability

The code used for super-resolution microscopy analysis is available on GitHub (https://github.com/super-resolution/Eiring-et-al-2023-supplement).

## Methods

### Fluorescence labeling

For microscopy measurements Cellvis chamber slides (8 well Chambered Coverglass Sytem #1.5 High Performance Cover Glass (0.17±0.005 μm), Cellvis) were coated with 0.1 mg/mL poly-D-lysine (PDL) for HEK293T cells or left untreated. Afterwards, cells were seeded into the chambers and allowed to adhere for one day. For ACE2 overexpression HEK293T cells were transfected for 24 h with human ACE2 (pCG1-hACE2, 200 ng DNA/well) using JetPrime (Promega) according to the manufacturer’s instructions. Then cells were fixed with 4 % methanol-free formaldehyde in HBSS for 10 min prior to staining with either 4 μg/ml anti-ACE2 (Biolegend, clone A20069I) or anti-NRP-1 (Biolegend, clone 12C2) antibodies for 1h. As a last step cells were fixed with 4% methanol-free formaldehyde / 0.2% glutaraldehyde before being imaged. All antibodies were labeled with AF647-NHS (Thermofisher, A20006) at a degree of labeling (DOL) of 2-3 to ensure specific binding and optimal imaging conditions.

### *d*STORM imaging

*d*STORM measurements were performed using an IX-71 inverted microscope equipped with an APON 60XOTIRF oil-immersion objective and an IX2-NPS nosepiece stage (all from Olympus, Hamburg, Germany)^47^. AF647 was excited with an appropriate laser system (Genesis MX 639 from Coherent, Göttingen, Germany). The excitation light was spectrally cleaned by appropriate bandpass filters and focused onto the backfocal plane of the objective. To switch between different illumination modes (Epi and TIRF illumination) the lens system and mirror were arranged on a linear translation stage. A polychromatic mirror (ZT405/514/635rpc, Chroma) was used to separate excitation (laser) and emitted (fluorescence) light. The fluorescence emission was collected by the same objective and transmitted by the dichroic beam splitter and detection filter (HC 679/41, Semrock), before being projected on an electron-multiplying CCD camera (iXon Ultra 897, Andor, Belfast, UK). The final pixel size of 128 nm was generated by placing additional lenses in the detection path. Excitation intensity was about ~3 kW/cm^2^. Typically, 15,000 frames were recorded at a frame rate of ~50 Hz (20 ms exposure time). To induce photoswitching of AF647 a PBS based buffer (pH 7.4) containing 100 mM β-mercaptoethylamin (Sigma-Aldrich) was used.

### Image reconstruction and data analysis

The recorded *d*STORM images were reconstructed with rapidSTORM 3.3 ^48^. Localization data acquired in *d*STORM measurements were filtered to remove background noise with less than 800 photons and analyzed with Locan^30^. For analysis of each *d*STORM image an appropriate region of interest at the basal membrane of the cell, was chosen. For clustering analysis, a DBSCAN clustering algorithm was applied to group detected localizations^49^. Suitable parameters were ε = 20 and minPoints = 3. Using these parameters allowed for quantification of detected localization within a certain distance giving ultimately information about existing oligomeric states and addressable receptors on the cell surface. We calculated and displayed Ripley’s H-function, a normalized Ripley’s K-function as previously described^50^. Computation was carried out for each ROI without edge correction. The averaged H-function was compared to those from 100 simulated data sets with localizations distributed on the same ROIs (and identical number of localizations in each ROI) according to complete spatial randomness or a Neyman-Scott process. The Neyman-Scott clustering process has homogeneously distributed parent events with each parent having n offspring events, where n is geometrically-distributed with mean equal to 9 (the average number of localizations recorded for a single antibody), and with the offspring positions having a Gaussian offset with a standard deviation of 10 nm. The maximum of the H-function indicates a distance that is between cluster radius and diameter and thus provides an estimate for the average cluster size.

### Pseudotyping of VSV with SARS-CoV-2 Spike protein (VSV-S^GFP^ production)

Pseudotyping of vesicular stomatitis virus (VSV) with the SARS-CoV-2 Spike protein was performed as previously described (Hoffmann et al., 2020 PMID: 32142651). Briefly, VSVΔG-GFP/luciferase was pseudotyped on BHK cells inducibly expressing the VSV G protein (kindly provided by Gert Zimmer) to generate VSVΔG-G. Next, 293T cells were transfected with a plasmid expressing SARS-CoV-2 Spike (kindly provided by Markus Hoffmann and Stefan Pöhlmann Hoffmann et al., 2020 PMID: 32142651) using TransIT-X2 (Mirus) according to the manufacturer’s instructions. At 24 h posttransfection, cells were infected with VSVΔG-G. In order to neutralize residual input virus, inoculated cells were washed twice with phosphate-buffered saline (PBS) 2 h postinfection, and new medium containing 1:1,000 anti-VSV-G antibody (8G5F11; Kerafast) was added. Supernatant containing replication-deficient VSVΔG pseudotyped with SARS-CoV-2 Spike protein (VSVΔG-S) was harvested 24 h postinfection, clarified by centrifugation, and stored at −80°C. Viral titers were determined on Vero E6 cells by quantifying GFP positive cells.

### Infection assay

To determine the percentage of infected cells, cells were seeded as described for *d*STORM analysis. On the next day cells were incubated with 250 μl VSV-S^GFP^/well for 24 h, washed with HBSS and subsequently fixed with 4% methanol-free formaldehyde / 0.1% glutaraldehyde in HBSS for 10 min. After washing with HBSS, nuclei were stained with NucBlue™ Fixed Cell ReadyProbes™ Reagenz (DAPI) (ThermoFisher, R37606). After an additional washing step with HBSS, the sample was analyzed on the EVOS™ FL Auto 2 microscope (Invitrogen). Each well was imaged automatically using a 20x objective and DAPI and GFP cubes to obtain two color images. In each image the nuclei and GFP expressing cells were counted using the particle analyzer in Fiji after applying unsharp masking and a threshold of 200. The percentage of infected cells was calculated for one well from all single images belonging to that well. The mean infectivity of a cell line was calculated from several wells in multiple experiments. Overall 1,144,292 (Vero), 747,301 (Vero E6), 1,053,538 (HEK293T), 729,943 (HEK293T:hACE2), 300,252 (COS-7) and 229,619 (U2-OS) cells were analyzed.

### GM1 staining

The glycolipid GM1 was stained with cholera toxin subunit B conjugated to AF647. Cells were seeded as described for *d*STORM analysis. The next day the sample was put on ice for 5 minutes, then stained with 5 μg/ml cholera toxin AF647 on ice for 20 min. After washing once with HBSS, cells were fixed with 4% formaldehyde / 0.2 % glutaraldehyde in HBSS for 10 min and washed again. Widefield imaging was performed on the ELYRA7 system (Zeiss, Germany) with a 25x objective.

### Reproducibility

All experiments were performed at least three times. Representative images are shown for each experiment.

### Reporting Summary

Further information on research design is available in the Nature Research Reporting Summary linked to this article.

---

**Extended Data Fig. 1.**
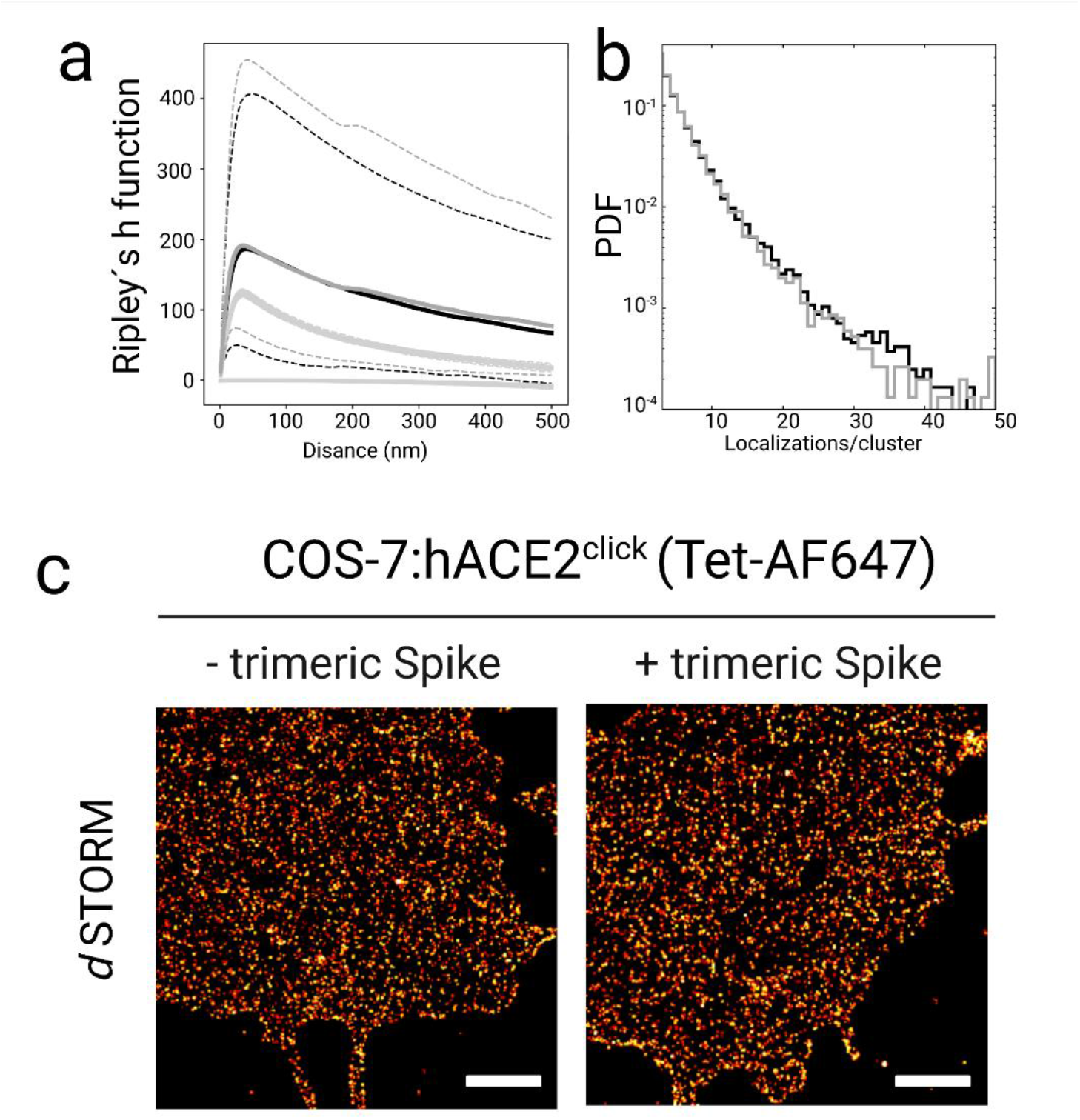
Binding of trimeric spike protein does not induce dimerization or trimerization of ACE2. **a,** Spatial distribution of localizations as analyzed by Ripley’s h function for ACE2^N137TAG^ (black) and ACE2^N137TAG^ in the presence of spike (gray) reveals identical clustering solely due to the repeated *d*STORM blinking events. For comparison, Ripley’s H function from 100 replicates of simulated data with spatial distributions following complete spatial randomness (bottom grey lines) or a clustered Neyman-Scott process (center grey lines) in identical ROIs are displayed (lower gray). Dotted lines indicate 95% confidence intervals of Ripley functions computed from 7 and 4 recordings, respectively. **b**, PDFs of localization numbers detected per localization cluster (as identified by DBSCAN yielding >3000 clusters for each condition) are identical confirming that ACE2 is present as monomer in the plasma membrane (black) and does not oligomerize in the presence of trimeric spike protein (grey). **c**, *d*STORM images of COS-7 cells expressing ACE2^N137TAG^ labeled with Pyr-Tet-AF647. Cells were either directly fixed after click labeling of first treated with recombinant trimeric spike protein (10 μg/ml for 20 min) before fixation and imaging. Scale bars, 1 μm (a).

**Extended Data Fig. 2.**
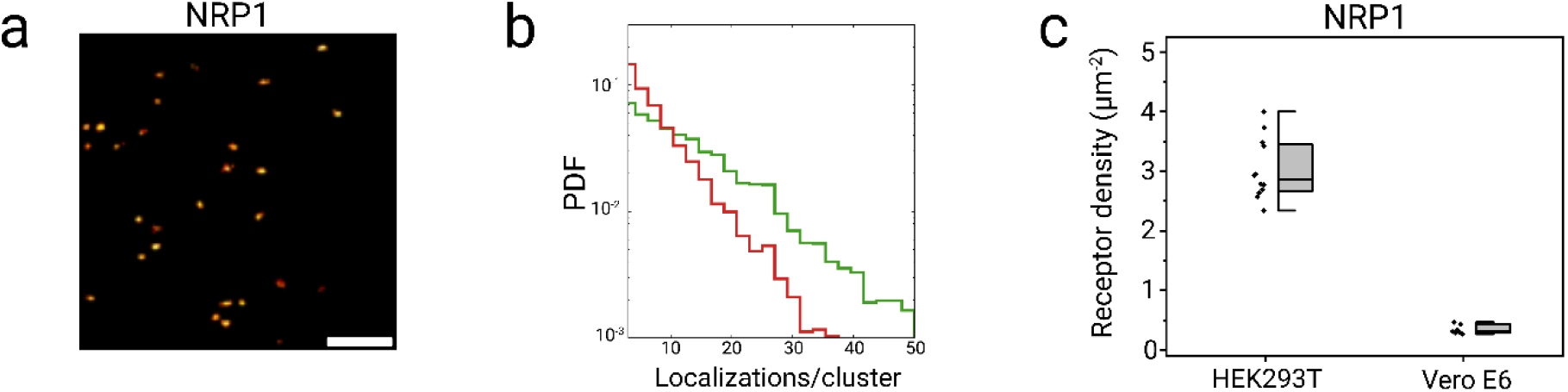
NRP-1 is identified as multimer in the membrane by *d*STORM. **a**, *d*STORM image of a HEK293T cell immunostained with AF647 labeled anti-NRP-1 primary antibody. **b**, PDF of localization numbers detected per NRP-1 localization cluster (green) showing a distribution typical for a dimer compared to monomeric ACE-2 (red). **c**, Boxplots NRP-1 expression determined from *d*STORM images with HEK293T, and Vero E6 cells. Vero cells show negligible expression of NRP-1. Scale bar, 3 μm (a).

**Extended Data Fig. 3.**
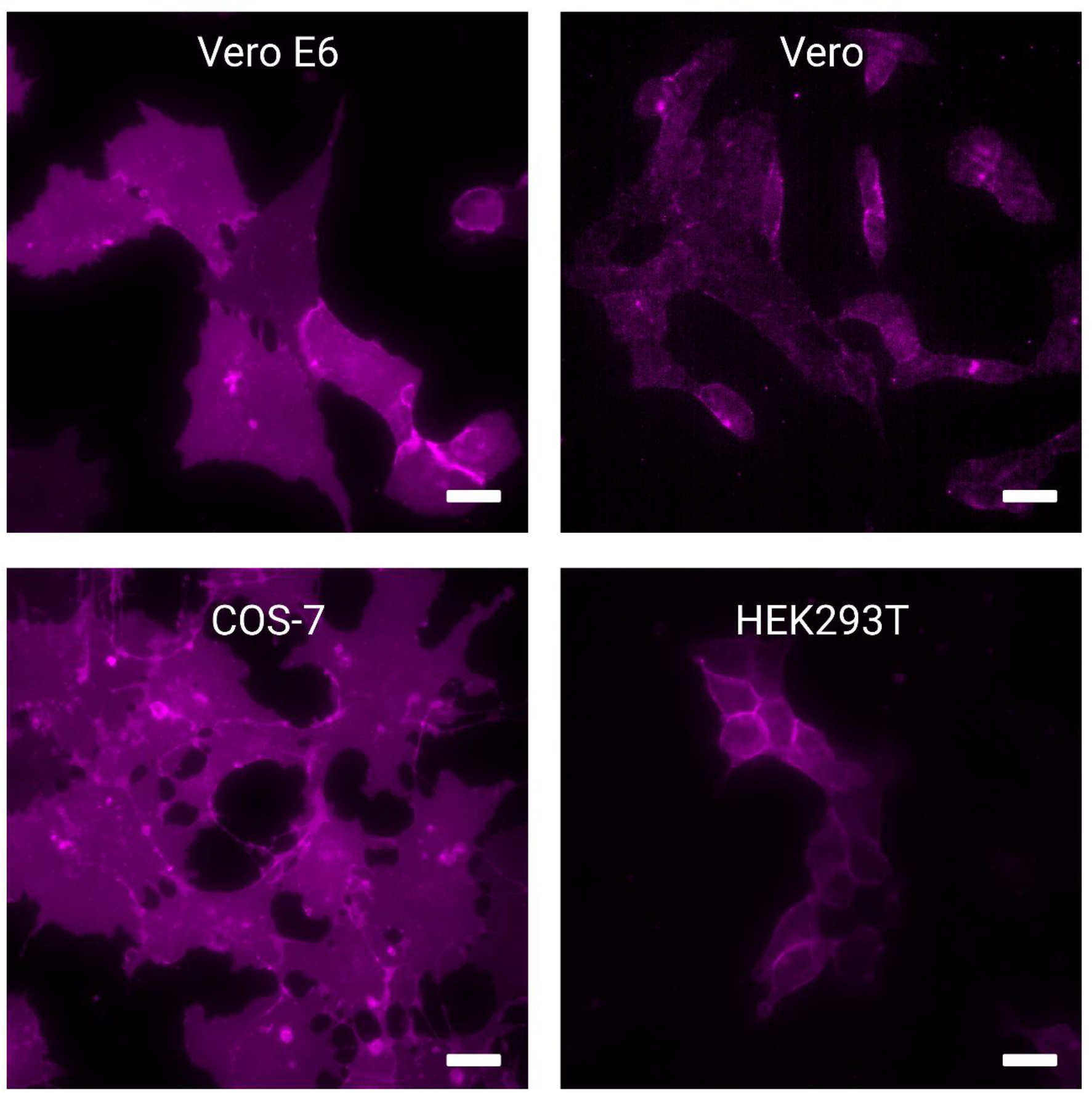
Higher GM1 concentrations in the plasma membrane support SARS-CoV-2 infection. Widefield fluorescence images of GM1 glycolipid abundance in different cell lines. GM1 was stained with 5 μg/ml AF647 labeled cholera toxin. Vero cells have substantially less GM1 present on the plasma membrane than Vero E6, HEK293T and COS-7 cells. Scale bars, 20 μm.

## Notes

https://github.com/super-resolution/Eiring-et-al-2023-supplement

### Competing Interest Statement

The authors have declared no competing interest.

