## Supplementary Information for "Coronaviruses use ACE2 monomers as entry receptors"

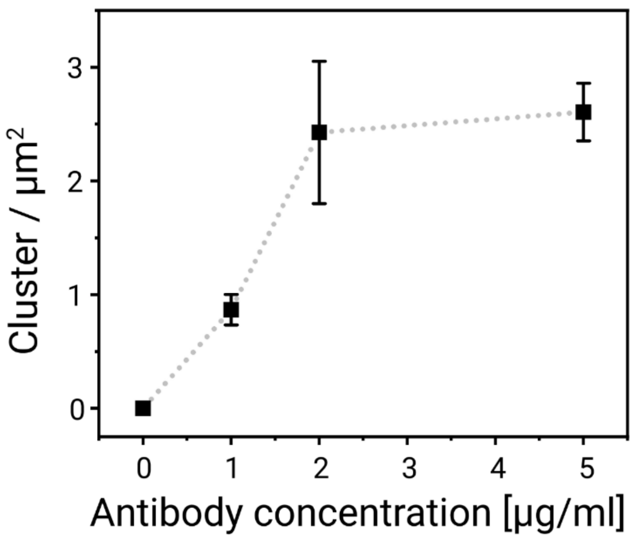


**Supplementary Fig. 1**. Average number of ACE2 localization clusters detected on the plasma membrane of Vero E6 cells at different antibody concentration. The data show that an antibody concentration between 2 and 5 µg ml$^{-1}$ is sufficient to label all accessible ACE2 epitopes in the plasma membrane.

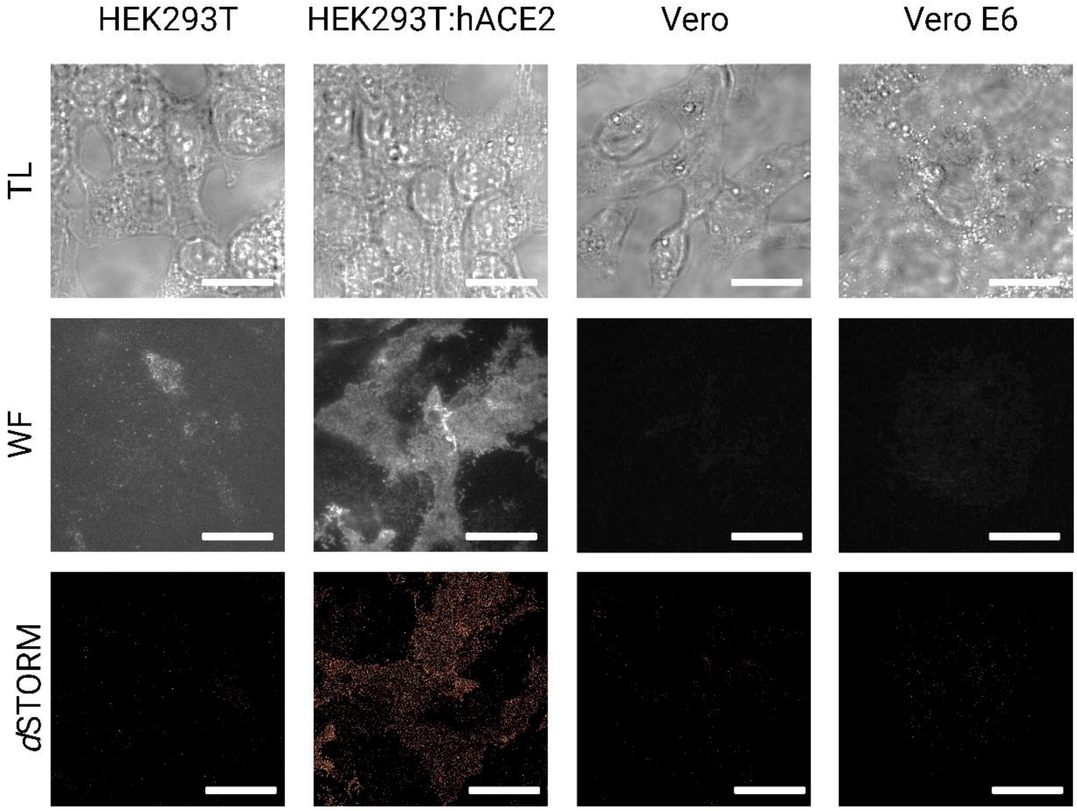


**Supplementary Fig. 2**. Transmitted light (TL), widefield fluorescence (WF) and *d*STORM example images of HEK293T, ACE2-overexpressing HEK293T (HEK293T:hACE2), Vero, and Vero E6 cells indicating different expression levels of ACE2. Scale bars, 20 µm.

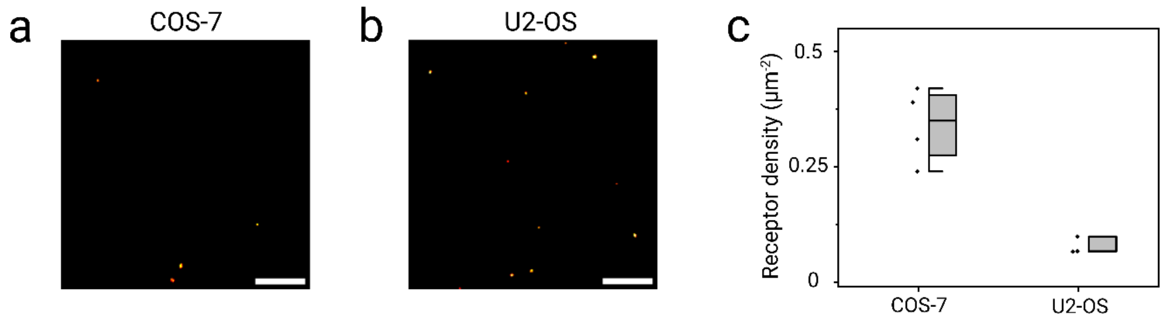


**Supplementary Fig. 3**. **a,b**, *d*STORM images and quantification of ACE2 on COS-7 and U2-OS cells lines. **c**, Boxplots show localization cluster densities translated into ACE2 receptors $\mu m^{-2}$ well below 0.5 demonstrating that both cell lines express ACE2 at negligible levels.

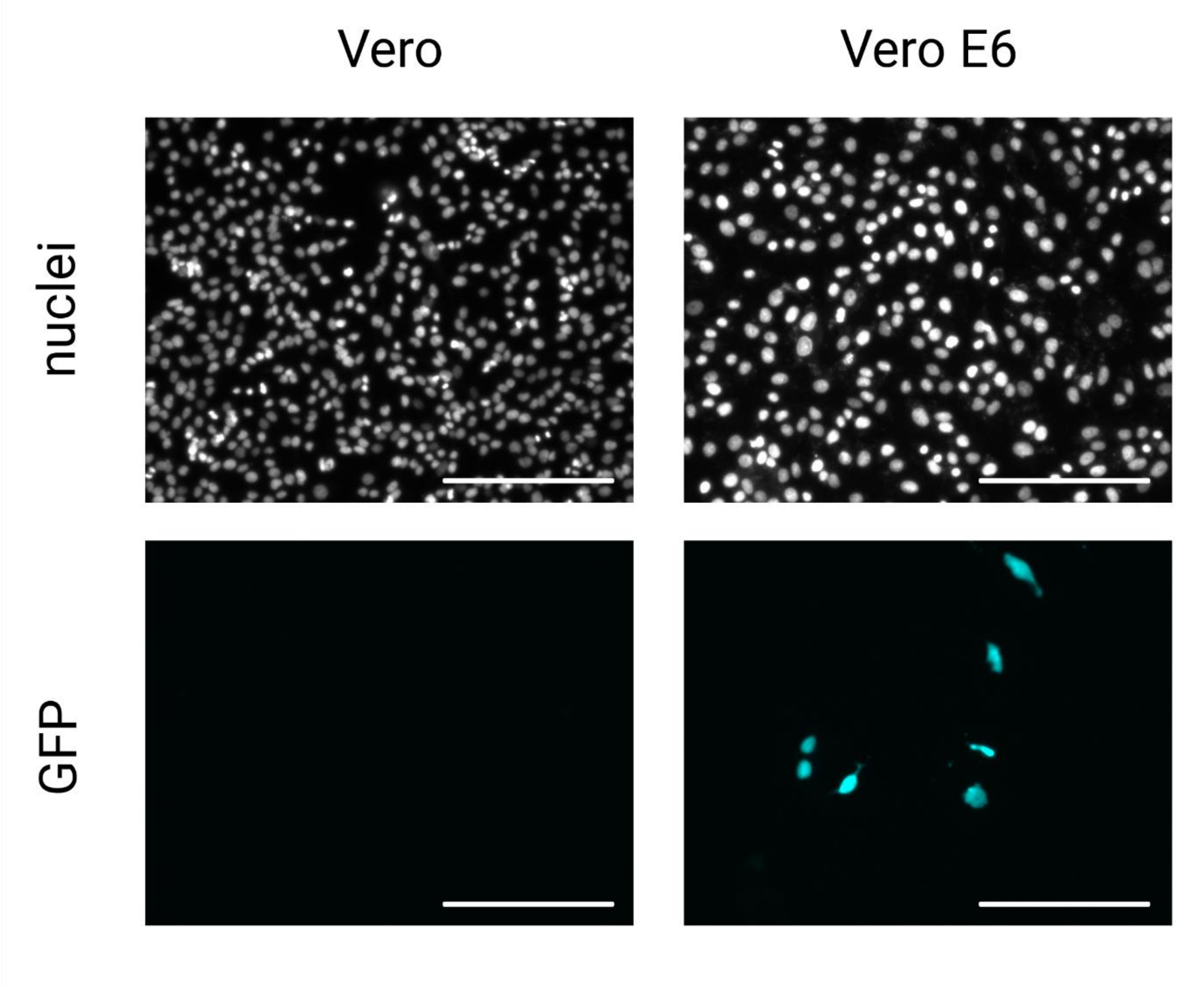


**Supplementary Fig. 4.** Widefield fluorescence images of DAPI (nuclei) and GFP in Vero and Vero E6 cells after 24 h infection with VSV-S$^{GFP}$. Scale bars, 200 µm.

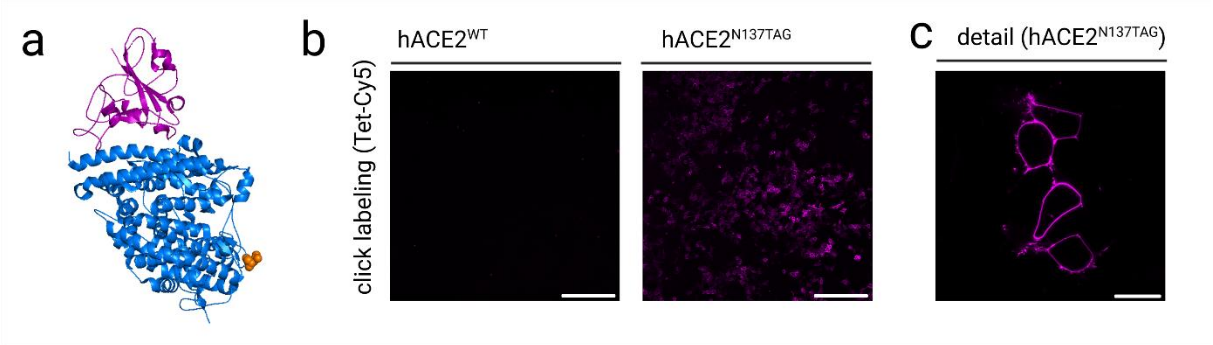


**Supplementary Fig. 5. a,** Crystal structure of the extracellular part of ACE2 (blue) and the receptor-binding domain (RBD) of the S protein of SARS-CoV-2 (purple). The asparagine (N) on position 137 (hACE2$^{N137TAG}$), which is substituted with an amber stop codon, is highlighted with orange spheres. Visualized with PyMOL. PDB Code: 2AJF. **b,** Click labeling of hACE2-WT (control) and hACE$^{N137TAG}$. Amber suppression was performed via co-expression of the previously described PylRS/tRNA$^{Pyl}$ pair[1] and addition of TCO*A-lysine. **c**, Detail magnified confocal fluorescence image of (b). Scale bars, 300 µm (b) and 20 µm (c).
